# maxnodf: an R package for fair and fast comparisons of nestedness between networks

**DOI:** 10.1101/2020.03.20.000612

**Authors:** Christoph Hoeppke, Benno I. Simmons

## Abstract

1. Nestedness is a widespread pattern in mutualistic networks that has high ecological and evolutionary importance due to its role in enhancing species persistence and community stability. Nestedness measures tend to be correlated with fundamental properties of networks, such as size and connectance, and so nestedness values must be normalised to enable fair comparisons between different ecological communities. Current approaches, such as using null-corrected nestedness values and *z*-scores, suffer from extensive statistical issues. Thus a new approach called NODF_*c*_ was recently proposed, where nestedness is expressed relative to network size, connectance and the maximum nestedness that could be achieved in a particular network. While this approach is demonstrably effective in overcoming the issues of collinearity with basic network properties, it is computationally intensive to calculate, and current approaches are too slow to be practical for many types of analysis, or for analysing large networks.
2. We developed three highly-optimised algorithms, based on greedy, hillclimbing and simulated annealing approaches, for calculation of NODF_*c*_, spread along a speed-quality continuum. Users thus have the choice between a fast algorithm with a less accurate estimate, a slower algorithm with a more accurate estimate, and an intermediate option.
3. We outline the package, and its implementation, as well as provide comparative performance benchmarking and two example analyses. We show that `maxnodf` enables speed increases of hundreds of times faster than existing approaches, with large networks seeing the biggest improvements. For example, for a large network with 3000 links, computation time was reduced from 50 minutes using the fastest existing algorithm to 11 seconds using `maxnodf`.
4. `maxnodf` makes correctly-normalised nestedness measures feasible for complex analyses of even large networks. Analyses that would previously take weeks to complete can now be finished in hours or even seconds. Given evidence that correctly normalising nestedness values can significantly change the conclusions of ecological studies, we believe this package will usher in necessary widespread use of appropriate comparative nestedness statistics.

## Introduction

Nestedness is a widespread and important feature of species interaction networks (Bascompte et al. 2003). Nestedness refers to the tendency for specialist species to interact with subsets of the species that more generalist species interact with. The prevalence of nested architectures, coupled with their high ecological and evolutionary importance, has given nestedness research a high profile, particularly for networks representing mutualistic interactions between species (Bastolla et al. 2009, Thébault & Fontaine 2010, James et al. 2012, Saavedra & Stouffer 2013, Suweis et al. 2013).

Like many indices of network structure, however, nestedness is correlated with other network properties, like connectance and the number of species in the network, which are themselves also highly correlated (Song et al. 2017, Ulrich et al. 2009). Additionally, many nestedness measures have bounds that are unconstrained to other network properties. For example, NODF, a popular measure of nestedness, is bounded between 0 and 1. This is problematic because fundamental constraints resulting from the size of the network and the number of links mean that, for many networks, maximum NODF values may be substantially less than 1 (Song et al. 2017). Therefore, comparing the nestedness of different networks using raw nestedness values should be avoided. Instead, it is essential to use nestedness metrics which are independent from network size, connectance and maximum nestedness (Song et al. 2017).

To resolve some of these issues, studies typically express nestedness values relative to a null expectation (for example, Welti & Joern 2015). Specifically, nestedness is expressed as a *z*-score: *z* = (Nestedness − *µ*)*/σ*, where *µ* and *σ* are the mean and standard deviation, respectively, of the nestedness values across an ensemble of networks generated using a particular null model. Problematically, it was recently shown that this method suffers from irrevocable statistical and inconsistency issues (Song et al. 2017), prompting the search for an appropriate way to compare the nestedness of different networks. Song et al. (2017) proposed a new normalised nestedness metric, NODF_*c*_, based on the NODF measure: NODF_*c*_ = NODF_*n*_ */*(*C*· *log*(*S*)), where NODF_*n*_ = NODF */max*(NODF), *C* is connectance, *S* is the geometric mean of the number of species in each level of the network (such as plants and pollinators or plants and seed dispersers), NODF is the raw NODF value for the network and *max*(NODF) is the maximum nestedness of a network with the same number of species and links as the focal network, subject to the constraint that every species has at least one link (Song et al. 2017). This new metric does not suffer from the statistical issues associated with *z*-scores and is thus robust for nestedness comparisons between networks (Song et al. 2017). To demonstrate this, Song et al. (2017) considered the long-standing prediction that networks are more ordered in less predictable environments (Levins 1968, May 1975). Previous studies using raw or *z*-score normalised nestedness values failed to find unified answers to this question, but by employing the NODF_*c*_ metric, Song et al. (2017) were able to confirm a positive associated between nestedness and temperature seasonality. This ability of the NODF_*c*_ metric to uncover patterns that previously could not be found is one of the strongest arguments for why it should be widely adopted.

While NODF_*c*_ is demonstrably a good statistic for nestedness comparisons, more technically, calculating the *max*(NODF) term in its formula is an NP-hard problem; the true maximum nestedness cannot be found in polynomial time. Instead, heuristic algorithms are required, that are not guaranteed to find the true optimum, but should at least find solutions close to the true optimum. Widespread adoption of the NODF_*c*_ approach is therefore highly dependent on the availability of fast algorithms that can find good solutions for the maximum nestedness of a network.

To date, two algorithms for this problem have been proposed. The first was a greedy algorithm by Song et al. (2017) (the ‘Song algorithm’) that was intuitive and achieved good optima, but was slow when run on large networks (33 minutes for a network with 797 species and 2933 interactions) (Simmons et al. 2019). This algorithm was refined by Simmons et al. (2019), who combined the greedy approach with simulated annealing (the ‘Simmons old’ algorithm). This new algorithm found higher levels of maximum nestedness than the Song algorithm, while reducing computation time by two thirds for large networks (11 minutes for the 797 species, 2933 interaction network). However, it was not available in `R`, which might limit its use among ecologists, and it was still too slow to be viable for many common types of analysis. For example, permutational approaches are frequently used in network ecology and would require calculating NODF_*c*_ hundreds or thousands of times. If the Song algorithm was used 1000 times on the large network mentioned above, the NODF_*c*_ calculations would take 22.9 days. While the ‘Simmons old’ algorithm reduces this to 7.6 days, this remains impractical. Even if permutational approaches are not necessary, run times of 33 minutes and 11 minutes, for the original and refined algorithms respectively, for a single large network are likely to deter users. Furthermore, if even larger networks are considered, such as the largest network in the Web of Life (www.web-of-life.es) database which has 1500 species and 15255 interactions, neither of these algorithms are likely to be practical for even a single calculation.

Thus while the normalised nestedness metric proposed by Song et al. (2017) is conceptually very robust, to date either slow implementations or suboptimal maxima has made the method impractical for all but simple analyses. However, it is essential that nestedness values are normalised correctly in order to ensure studies make accurate inferences. Here we fill this gap by introducing, `maxnodf`, an `R` package that enables rapid evaluation of NODF_*c*_, for the first time making correctly-normalised nestedness values accessible for even complex analyses of large networks. Below we describe the package and its implementation, alongside comparative performance benchmarking and two examples of how it can be applied to empirical data.

We note that while our focus here is on species interaction networks, nestedness is also found in patterns of species occurrence. `maxnodf` is equally applicable to these data.

## Description

### Improving NODF calculation efficiency

The NODF_*c*_ metric relies on evaluating 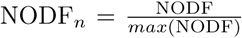, where *max*(NODF) denotes the maximum nestedness that can be obtained by networks in the same class as the original network. We say that two network are in the same class if they share the same number of species in both classes and the total number of links in the network are identical.

We can compute the nestedness of a single network with reasonable efficiency, however approximating *max*(NODF) requires the application of optimisation algorithms. During the course of such algorithms, it is often the case that NODF is reevaluated repeatedly. The speed at which we can reevaluate NODF thus becomes the determining factor for the performance of optimisation algorithms.

To assess the computational cost of NODF, we revisit the definition provided in Almeida-Neto et al. (2008) and provide an algorithm for its evaluation. The computation of NODF relies on evaluating, for all row and column pairs, the paired nestedness 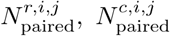. We consider the rows *r*_1_, *r*_2_ ∈ ℕ_≥1_. If *r*_2_ ≥ *r*_1_ we set 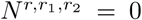. Next we check that the number of links from row *r*_1_ to row *r*_2_ is decreasing. If the number of links is non decreasing, we set again 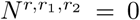, otherwise we compute the partial overlap between rows *r*_1_ and *r*_2_. The partial overlap is defined as the proportion of links in row *r*_2_, which have matching links in row *r*_1_. The paired nestedness for columns is defined analogous. For a network *A* ∈ {0, 1}^*N*×*M*^ with *N* rows and *M* columns, computing the partial overlap between two rows or columns requires 2*M* or 2*N* operations respectively. During the computation of NODF we need to compute *N*^2^ paired nestedness values for rows and *M*^2^ paired nestedness values for columns. This results in a computational cost of *𝒪* (*N*^2^ × *M* + *M*^2^ × *N*) for one NODF evaluation. This cost can also be seen in Algorithm 1, as it results from summation inside of nested for loops. As a result, evaluating NODF for a network of twice the size *Ã* ∈ {0, 1} ^2*N*×2*M*^ requires 8 times as many operations as evaluating NODF for the original network *A*. We thus say that the computational cost of NODF grows cubic with network size.

#### Algorithm 1 NODF evaluation

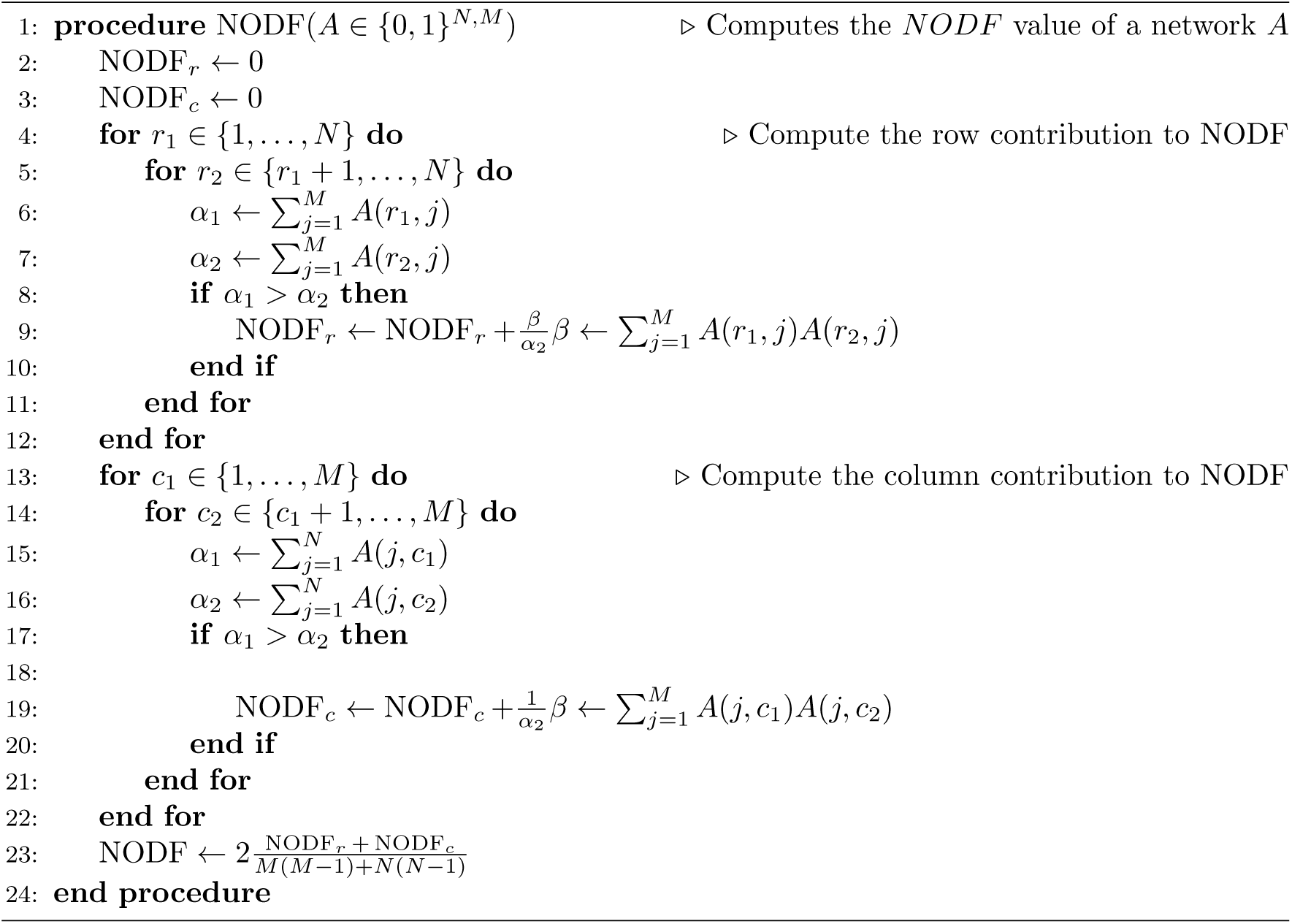

During optimisation processes we often perform minor modifications, which only alter the network in a single position. We can use the knowledge that all but one entries in the network remain unchanged to accelerate the computation of NODF. Consider a modification of network *A* ∈ {0, 1}^*N* ×*M*^ in position (*i, j*). We only need to consider the *N* values *A*(1, *j*), *… A*(*N, j*) to update NODF_*r*_ and the *M* values *A*(*i*, 1), *…, A*(*i, M*) to update NODF_*c*_. This results in a total complexity reduction from *𝒪* (*N*^2^ × *M* + *N* × *M*^2^) to *𝒪* (*N* + *M*). We can thus recompute the NODF value for a network which has been modified in a single position in linear rather than cubic time. These efficiency improvements enable the use of more advanced algorithms like, like hill climbing or simulated annealing, to the NODF maximisation problem.

### Package functions

The *R* package `maxnodf` contains methods for maximising the NODF metric for ecological networks. Different applications will require a different trade-off between optimisation quality and performance. To address these different preferences we offer three different quality settings in the `maxnodf` procedure. These quality settings build upon each other, guaranteeing that the NODF values obtained on a higher quality setting will never be below those obtained at lower quality.

We note that networks with *N* rows *M* columns and a total of *L* links, where *L* ≤ *N* + *M* are necessarily compartmentalised (contain multiple disconnected subnetworks). In such cases nestedness computations should be performed on individual compartments of the network. In these cases `maxnodf` displays a warning message, and the maximisation is instead performed on the set of networks with *L* = *N* + *M* + 1 links.

### Quality 0: Greedy algorithm

Calling `maxnodf` with a quality parameter of 0 invokes a greedy optimiser, which is the least accurate but fastest optimiser offered in `maxnodf`. We first construct a network *A*^(0)^ ∈ {0, 1}^*N*×*M*^ with *A*^(0)^(1, *k*) = *A*^(0)^(*j*, 1) = *A*^(0)^(2, 2) = 1 ∀*j* = 1, …, *N, k* = 1, …, *M*. For the case *N* = *M* = 5 the initial network is given by

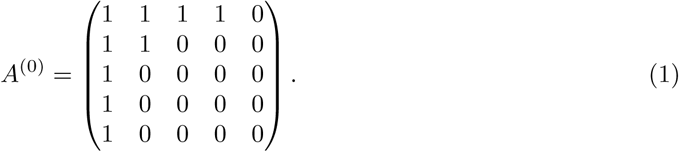

We denote the network and its component at step *k* by *A*^(*k*)^ and 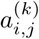 respectively. In each step we consider potential new link positions, which border links in the current network {(*i, j*)|*a*^(*k*−1)^(*i, j*) = 0, *a*^(*k*−1)^(*i* − 1, *j*) = *a*^(*k*−1)^(*i, j* − 1) = 1}. We evaluate NODF for each of the potential modifications of *A*^(*k*−)^ and introduce a new link at the NODF maximising index pair (*i*\*, *j*\*). We successively add links to *A*^(*k*)^ until the desired number of links is reached.

### Quality 1: Greedy algorithm with hill climbing

Calling `maxnodf` with the quality parameter 1 first invokes the same greedy algorithm as quality 0 does. Starting further optimisation from this network ensures that quality 1 will never perform worse than quality 0.

Improvements over the greedy algorithm can be obtained with an algorithm called hill climbing. In hill climbing we perform local modifications to a network by moving a link from its original position to a neighboring position. For all link positions (*i, j*), with *i, j* ≥ 2 and *A*(*i, j*) = 1 we test moving the link to all possible neighbor positions, and for each of the resulting networks we compute NODF. In each hill climbing step, the network is modified by executing the NODF-maximising move of a link to a neighboring position. Links located in the first row and first column are excluded, to avoid compartmentalization effects during this optimisation procedure. The hill climbing algorithm terminates once the no such local modification achieves a strictly higher NODF value. This ensures that we always terminate on a local maximum.

### Quality 2: Greedy algorithm with hill climbing and simulated annealing

For the quality parameter of 2, we first apply the same greedy and hill climbing algorithms as in quality settings 0 and 1. This ensures both that quality 2 will never perform worse than 0 or 1 and that we start the NODF maximisation procedure from a local optimum. As a result, moving a link to neighboring position cannot improve the NODF value. To further improve on such networks we require algorithms that are able to escape from local optima. Such algorithms necessarily have the property that worse results need to be accepted in certain situations.

One such algorithm is called simulated annealing and was proposed by Kirkpatrick et al. (1983). In simulated annealing, we accept network modifications which reduce the NODF with a certain probability. This probability is determined by a temperature parameter *T* and the reduction in NODF which this move has introducedΔ NODF. We then accept the move according to the Kirkpatrik acceptance probability function

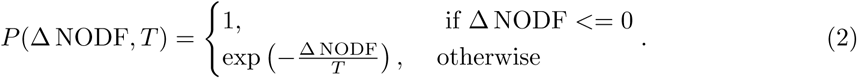

After each step of the simulated annealing algorithm, the temperature *T* is reduced by setting *T*^(*k*)^ = *αT*^(*k*−1)^, where 0 *< α <* 1. As result the algorithm is more likely to accept large drops in NODF when *T* is large, resulting in a fast scan across the space of possible networks at the start of the algorithm, followed by a more concentrated search in the area where we expect the true optimal network, when *T* is low. We combine the simulated annealing with hill climbing by starting a search for local optima whenever the simulated annealing algorithm reaches a new NODF-maximising network. This condition guarantees that the hill climbing algorithm does not reset the algorithm the previous local optimum. The algorithm used by `maxnodf` at quality 2 is given below in pseudocode.

#### Algorithm 2 SimulatedAnnealing

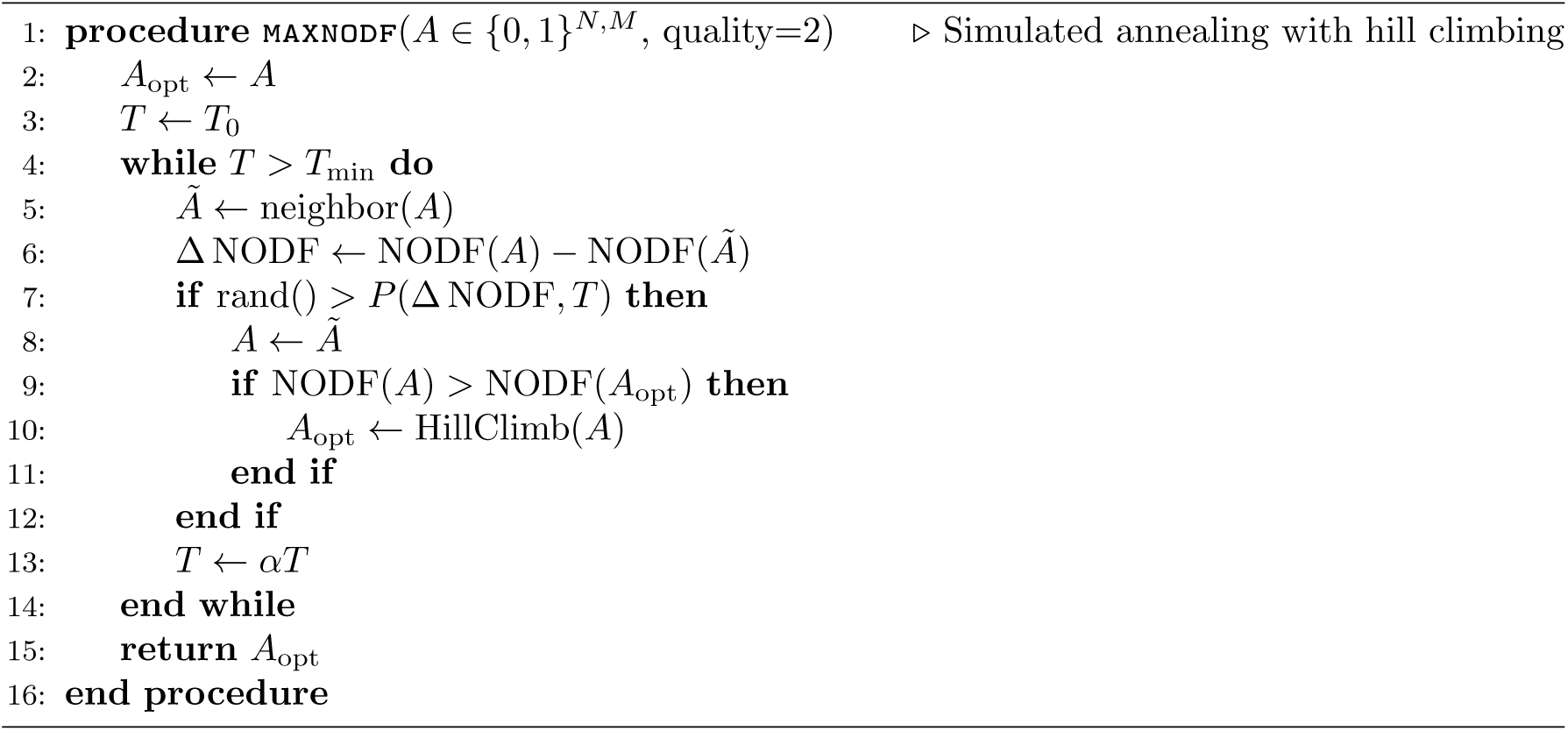

## Performance

We recorded the time to compute, and maximum NODF achieved, when calculating max(NODF) using five algorithms: the three algorithms in the `maxnodf` package (quality 0, 1 and 2), the original greedy algorithm proposed by Song et al. (2017) (‘Song’), and the refinement of this algorithm proposed by Simmons et al. (2019) (‘Simmons old’). Algorithms were run on all pollination networks from the Web of Life dataset (http://www.web-of-life.es), excluding 13 networks with either more than 3000 links or which were not a single, connected component. These inclusion criteria ensured all algorithms could be run on the entire dataset: the Song algorithm was impractically slow on networks with more than 3000 links, and the algorithms cannot be run on networks which comprise multiple, disconnected subnetworks. This resulted in a dataset of 135 networks. Timings were carried out on a computer with an i7-8550U (1.8 GHz) processor with 16 GB RAM (2133 MHz).

Overall, the variation in runtime was substantial, spanning five orders of magnitude. We find that, on average, the slowest algorithm is the ‘Simmons old’ algorithm, followed by the Quality 2 algorithm, the original Song algorithm, the Quality 1 algorithm and finally the Quality 0 algorithm, which is the fastest (Figure 1a). This ordering is broadly expected: simulated annealing algorithms, like ‘Simmons old’ and Quality 2, achieve better optima but are slower, while greedy algorithms, like Song and Quality 0, achieve worse optima but are faster. The `maxnodf` Quality 2 algorithm is the fastest simulated annealing algorithm, and the Quality 0 algorithm is the fastest greedy algorithm. Notably, the Quality 1 algorithm, which uses greedy and hillclimbing components, would be expected to take an intermediate time, slower than greedy algorithms, but faster than simulated annealing. However, instead we find that Quality 1 is actually faster on average than the Song greedy algorithm, while also achieving better optima (Figure 1a).

**Figure 1:**
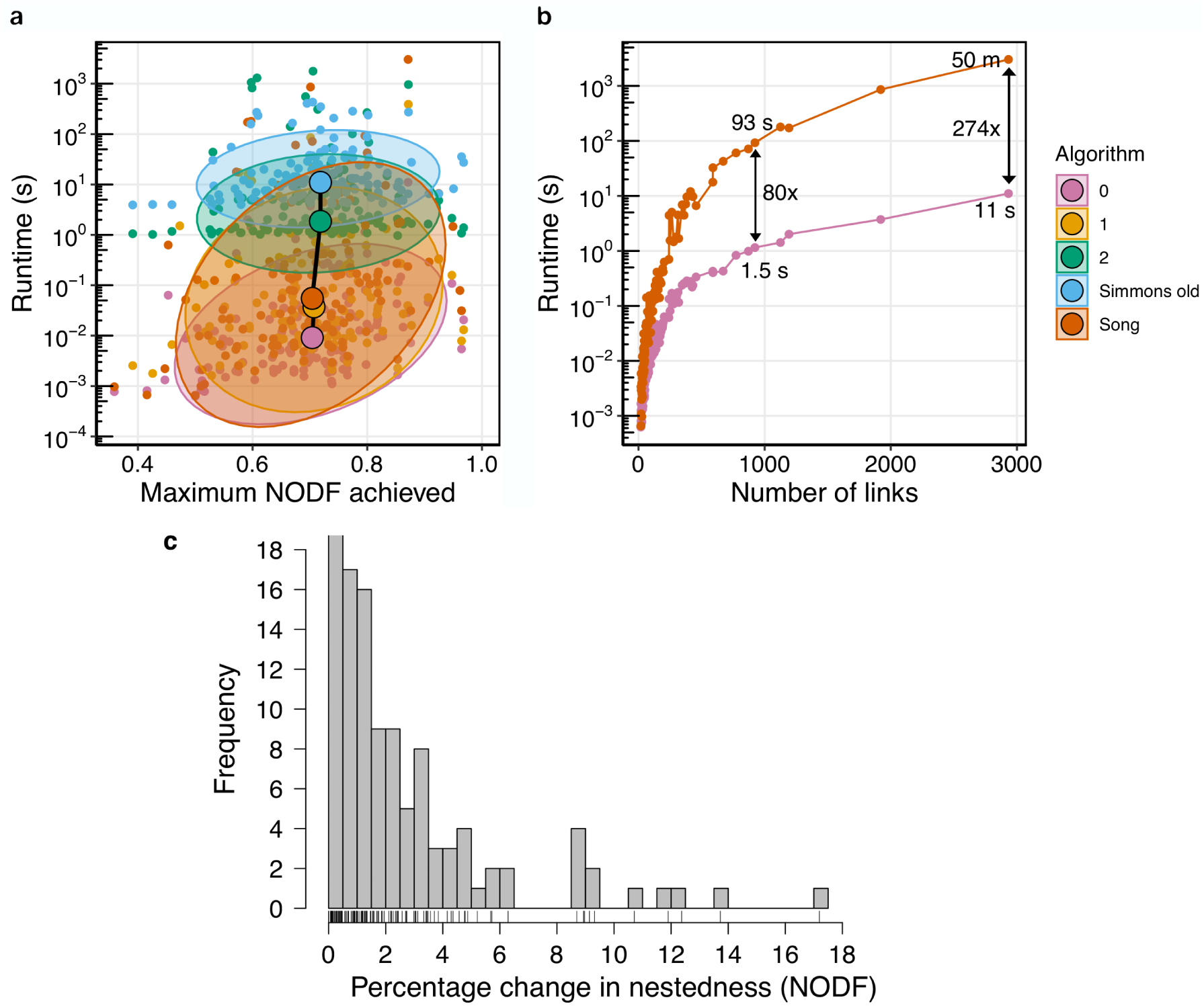
Comparative benchmarking of nestedness maximisation algorithms. **(a)** Comparison of the time taken to run, and maximum NODF achieved, for: (i) the three algorithms in the `maxnodf` package (quality 0, 1 and 2); (ii) the algorithm proposed by Simmons et al. (2019); and (iii) the original algorithm proposed by Song et al. (2017). All algorithms were run on an identical set of 135 empirical pollination networks from the Web of Life dataset. Small points represent individual networks; large points represent medians. Ellipses represent 95% confidence intervals. **(b)** Comparison of how long the quality 0 greedy algorithm and the original greedy algorithm proposed by Song et al. (2017) take to run on networks with different numbers of links. Network data were the same 135 networks as in (a). Arrows show the time difference between the two algorithms for particular networks. The label of the arrow shows how many times faster the quality 0 algorithm was (e.g. ‘80x’ is 80 times faster), while the numbers at the ends of the arrow show the time each algorithm took to complete for a particular network (e.g. ‘50m’ = 50 minutes, ‘11s’ = 11 seconds.) **(c)** Percentage increase in maximum NODF achieved by the Quality 2 algorithm compared to the Quality 0 algorithm for all 135 networks in the dataset.

The above discussion focuses on the average performance of each algorithm, but average values can mask important patterns. Given that the Quality 0 algorithm will be the most widely used, in Figure 1b, we compare its performance to that of the original Song algorithm. Note that these two algorithms are equivalent: the `maxnodf` version is simply a faster implementation of the Song algorithm. The speed improvement offered by our implementation is substantial, and is greatest for networks with larger numbers of links: for the largest network in our dataset, our algorithm offers a 274 times speed improvement, reducing the computation time from 50 minutes to 11 seconds (Figure 1b). For networks with fewer links, the improvement is still large, becoming increasing less important for the networks with the fewest links. These performance improvements enable complex analyses to use the *max*(NODF) approach, even for large networks, while this was unlikely to be possible previously.

In terms of maximum NODF achieved, as expected the ‘Simmons old’ and Quality 2 simulated annealing algorithms perform best, while the Song and Quality 0 greedy algorithms perform worst (Figure 1a). The magnitude of the improvement afforded by the slower algorithms is, however, generally small: on average, the Quality 2 algorithm produces maximum NODF values that are 2.3% higher than those produced by Quality 0. However, again, this average masks some variation, with improvements of up to 17% (Figure 1c). Thus while the Quality 0 algorithm will be suitable for most purposes, the higher-quality algorithms are available if only a small number of networks are being studied, or if the most accurate NODF values are needed.

## Applications

Here we conduct two short analyses. First, we use pollination networks from the Web of Life dataset (www.web-of-life.es) to determine the level of collinearity between raw NODF values and NODF_*c*_. Specifically, we calculate NODF and NODF_*c*_ for all networks and test for a correlation in their ranks. We find no correlation in ranks (Spearman’s: *ρ* = -0.11, S = 831880, *P* = 0.16; Figure 2a), indicating that the most nested network as measured by raw NODF is not the most nested as measured by NODF_*c*_ (and so on). This demonstrates the importance of normalising nestedness correctly — the relative nestedness of networks when normalised is very different to when they are not normalised. Second, we compare how nestedness varies over time when measured using NODF and NODF_*c*_. Data was from four plant-pollinator communities in the Seychelles, sampled for eight consecutive months from September 2012 to April 2013 (Kaiser-Bunbury et al. 2017). To measure the stability of nestedness over time, we calculated the coefficient of variation in raw and normalised nestedness for each of the four communities. We found that nestedness appears more stable over time when measured using NODF_*c*_ (t = 4.80, df = 3, *P* = 0.02), suggesting that the macro structure of these communities may be more stable than would be recognised from using raw NODF values.

**Figure 2:**
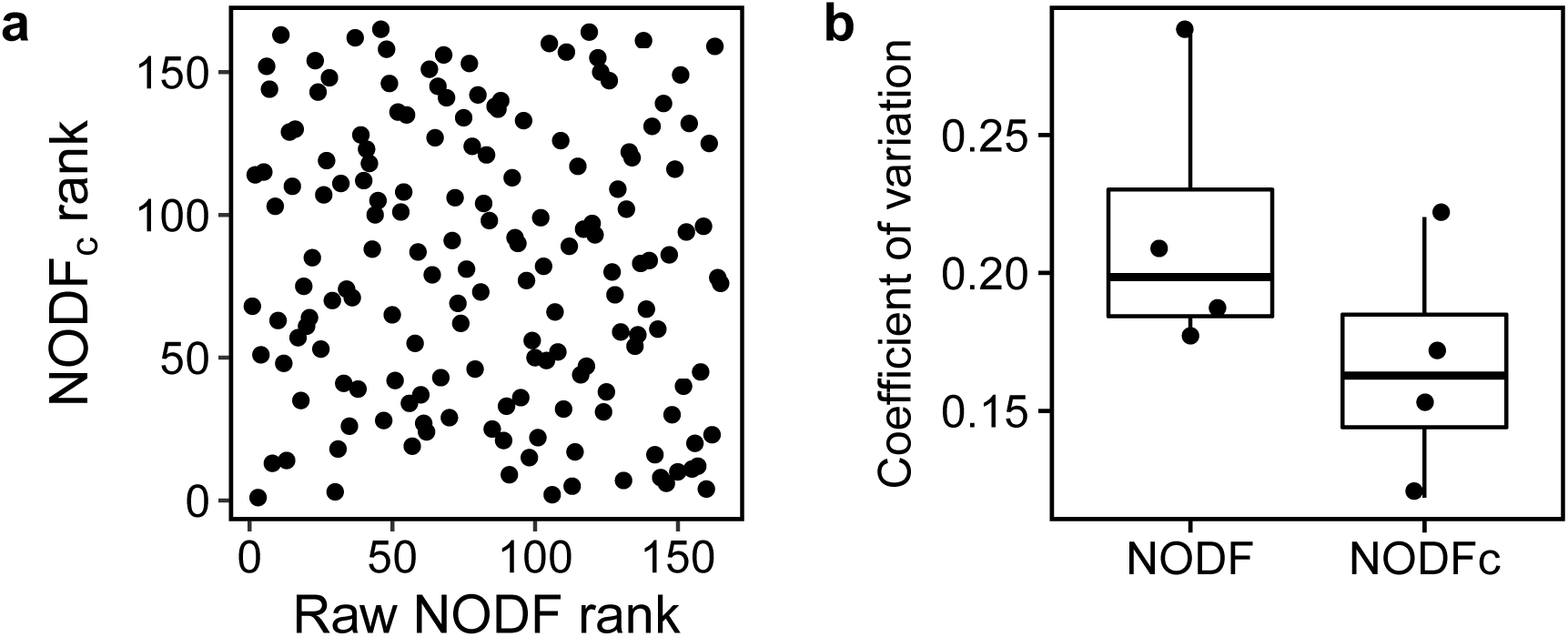
(a) The relationship between the ranks of nestedness values in raw and normalised forms. (b) Coefficient of variation of nestedness values in four networks over time, measured using raw NODF and NODF_*c*_. NODF_*c*_ values indicate that nestedness values are more stable over time than is shown with the raw nestedness values.

## Implementation and availability

The `maxnodf` package is available for the `R` programming language. To install the package, run `install.packages (“maxnodf”)`. This paper describes version 1.0.0 of the software. The source code of the package is available at https://github.com/christophhoeppke/maxnodf. Any problems can be reported using the Issues system. The code is version controlled with continuous integration and has code coverage of approximately 95%. All code is released under the MIT license.

## Conclusions

Nestedness is pervasive pattern in ecological systems. In particular, nestedness measures have been widely used in studies of mutualistic species interaction networks, but a lack of proper normalisation has limited our ability to make inferences. `maxnodf` is the first package to implement rapid calculation of the NODF_*c*_ metric, a nestedness measure that is demonstrably fair to compare between networks of different sizes and connectances. The package contains three optimised algorithms that allow the user to choose their own trade off between speed and quality. We anticipate that, by making NODF_*c*_ calculations feasible for complex analyses and for use with large networks, `maxnodf` will usher in widespread adoption of correctly-normalised nestedness values.

## Acknowledgements

CH and BIS thank the Cambridge Faculty of Mathematics CMP bursary fund for support. BIS was supported by the Natural Environment Research Council as part of the Cambridge Earth System Science NERC DTP (NE/L002507/1) and by a Royal Commission for the Exhibition of 1851 Research Fellowship. CH was supported by the Engineering and Physical Sciences Research Council EPSRC, the University of Oxford Clarendon fund, and the John Henry Jones fund of Balliol College Oxford.

## Author contributions

BIS conceived the study; CH and BIS developed the algorithms, conducted analyses and wrote the first manuscript draft.

## Data accessibility

All networks used in this study are available from the Web of Life repository (www.web-of-life.es)

